# An Arrayed Genome-Wide Perturbation Screen Identifies the Ribonucleoprotein hnRNP K As Rate-Limiting for Prion Propagation

**DOI:** 10.1101/2022.03.03.482765

**Authors:** Merve Avar, Daniel Heinzer, Alana M. Thackray, Yingjun Liu, Marian Hruska-Plochan, Stefano Sellitto, Elke Schaper, Daniel P. Pease, Jiang-An Yin, Asvin K.K. Lakkaraju, Marc Emmenegger, Marco Losa, Andra Chincisan, Simone Hornemann, Magdalini Polymenidou, Raymond Bujdoso, Adriano Aguzzi

## Abstract

A defining characteristic of mammalian prions is their capacity for self-sustained propagation. Theoretical considerations and experimental evidence suggest that prion propagation is modulated by cell-autonomous and non-autonomous modifiers. Using a novel quantitative phospholipase protection assay (QUIPPER) for high-throughput prion measurements, we performed an arrayed genome-wide RNA interference (RNAi) screen aimed at detecting modifiers of prion propagation. We exposed prion-infected cells in high-density microplates to 35’364 ternary pools of 52’746 siRNAs targeting 17’582 genes representing the mouse protein-coding transcriptome. We identified 1191 modulators of prion propagation. While 1151 of these modified the expression of both the pathological prion protein, PrP^Sc^, and its cellular counterpart PrP^C^, 40 genes affected selectively PrP^Sc^. Of the latter, 20 genes augmented prion production when suppressed. A prominent limiter of prion propagation was the heterogeneous nuclear ribonucleoprotein Hnrnpk. Psammaplysene A (PSA), which binds Hnrnpk, reduced prion levels in cultured cells and protected them from cytotoxicity. PSA also reduced prion levels in infected cerebellar organotypic slices and alleviated locomotor deficits in prion-infected *Drosophila melanogaster* expressing ovine PrP^C^. Hence, genome-wide QUIPPER-based perturbations can discover actionable cellular pathways involved in prion propagation. Finally, the unexpected identification of a prion-controlling ribonucleoprotein suggests a role for RNA in the generation of infectious prions.

## Introduction

The life cycle of mammalian prions entails the misfolding and aggregation of the cellular protein PrP^C^ and its incorporation into a nucleated higher-order isoform called PrP^Sc^ (Aguzzi and Calella, 2009). Once the PrP^Sc^ aggregates reach a critical size, they break and elongate again by recruiting additional monomers (Knowles et al., 2009).This cyclic sequence of events is the basis for the increase in prion infectivity (Nuvolone et al., 2009). However, it is still unknown whether this process occurs autonomously akin to crystal growth, or if it necessitates auxiliary cofactors (Deleault et al., 2012a; Deleault et al., 2012b). The latter is suggested by the observation that propagation of prions in a cell-free system is inefficient and necessitates extreme conditions such as cyclic high-energy sonication, shaking, or partial chemical denaturation (Atarashi et al., 2011; Saborio et al., 2001). In contrast, infection of animals or cultured cells with prions can yield titer increases by several orders of magnitude under physiological conditions (Klöhn et al., 2003; Prusiner et al., 1982). This suggests that living systems contain important cofactors that enable prion propagation, e.g., by lowering the thresholds of rate-limiting reactions.

How could one possibly identify such cofactors? In the case of other neurodegenerative diseases, crucial insights were derived from human genetics. The study of families afflicted by inherited forms of Alzheimer’s and Parkinson’s disease have yielded a plethora of genes encoding proteins directly linked to the offending aggregates (Jansen et al., 2019; Nalls et al., 2019; van Rheenen et al., 2016). However, this approach has not been as successful in the case of prion diseases, partly because of their rarity which precludes large genome-wide association studies (Lloyd et al., 2013). As a result, the only modifiers robustly associated with predisposition to prion diseases are genetic polymorphisms within the PrP^C^-encoding *PRNP* gene itself (Jones et al., 2020; Mead et al., 2009; Mead et al., 2012; Sanchez-Juan et al., 2014).

A possible approach to this conundrum consists of investigating candidate genes which may be inferred from existing reports or from their role in phenomena pertinent to prion propagation. For example, transcription factors involved in *PRNP* mRNA expression (Bellingham et al., 2009; Dery et al., 2013; Rybner et al., 2002; Vincent et al., 2009), or proteins involved in its degradation (Parkyn et al., 2008; Shyu et al., 2002; Vincent et al., 2009), may represent such candidates. However, this approach has major limitations. Any potential candidates, in order to be identified as such, must have been described previously in similar contexts. Consequently, any fundamentally novel mechanisms cannot be discovered because they would not exist as priors.

Forward genetic screens, in which each protein-coding gene is being modified and hits are identified by their effect on the phenotype of interest, represent a less biased and more inclusive approach with the potential of yielding wholly unpredicted hits. Moreover, the identification of relationships between hits, e.g., because they fall within a single pathway, or because they encode individual components of a single physical complex, can fortify the confidence in the validity of the results. In the past, such screens have been most effectively performed in unicellular organisms that undergo a haploid phase, such as yeast (Giorgini et al., 2005; Outeiro and Lindquist, 2003). However, more recent technologies such as RNA interference (RNAi) and CRISPR have enabled the deployment of forward genetic screens in diploid mammalian cells (Heinzer et al., 2021; Kampmann, 2018; Mohr et al., 2010).

In this work, we have used arrayed RNAi to interrogate all genes of the mouse genome for their influence on prion propagation. We have discovered 40 such genes. 20 of these were found to reduce prion propagation when suppressed, but 20 genes enhanced prion propagation when silenced. Some of these modifiers fell within pathways expected to control prion propagation (Marbiah et al., 2014). However, others were entirely surprising, including the small heteronuclear RNA binding protein, Hnrnpk.

## Results

### Establishment of a genome wide high-throughput screen for identification of prion modulators

Scalable, reproducible high-throughput assays should consist of only a few steps and should not require analyte transfers to different reaction containers. Immunochemical prion detection (by Western blotting, enzyme-linked immunoassay, or other methods) is typically preceded by limited proteolysis using proteinase K (PK), eliminates PrP^C^ and ensures that any residual signal is specific to PrP^Sc^ (Bolton et al., 1982). However, digestion with PK requires fastidious titration and accurate timing, which may introduce confounders (McKinley et al., 1983).

To solve these issues, we took advantage of phosphatidylinositol-specific phospholipase C (PIPLC), an enzyme that cleaves proteins attached to the membrane via a glycophosphatidylinositol (GPI) anchor from the surface of cells (Heinz et al., 1995). While PrP^C^ is GPI-anchored (Stahl et al., 1990), prion aggregates appear to associate in cells independently of the anchoring. PIPLC treatment of intact cells leads to the release of most of the PrP^C^ into the supernatant, while prions remain cell associated, (Borchelt et al., 1993; Stahl et al., 1990) and in endocytic compartments (Taraboulos et al., 1992).

Prion assemblies were disaggregated using sodium hydroxide (NaOH, pH=14, 66.6 mM) (Peretz et al., 2001), neutralized with NaH_2_PO_4_ buffer (pH=4.5, 83.3 mM) to near-neutral pH (Li, 2016), and a Förster energy transfer donor-acceptor antibody pair (Allophycocyanin-POM1 (APC-POM1), Europium-POM19 (EU-POM19)) was added (Ballmer *et al*., 2017; Pease *et al*., 2019; Polymenidou et al., 2008). PrP was then detected by time resolved (TR) FRET (Ballmer et al., 2017; Heinzer et al., 2021; Pease et al., 2019). We termed the resulting assay QUantItative Prion PhospholipasE pRotection assay (QUIPPER, Fig. 1A).

**Figure 1.**
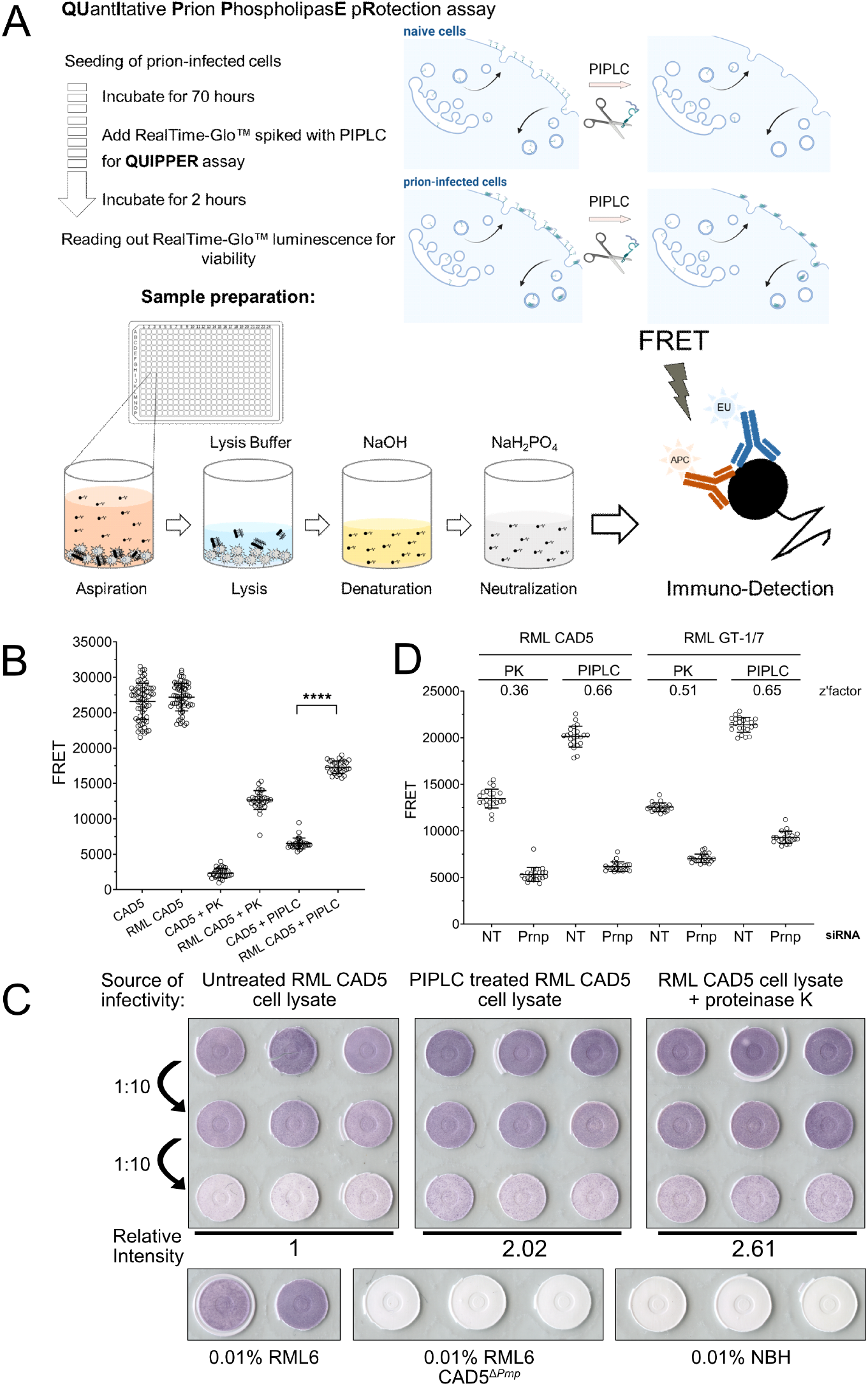
A cell-based high-throughput prion detection assay for an arrayed whole genome RNAi screen. **A:** Workflow of the QUIPPER assay. **B:** PK vs. PIPLC treatment to determine prion loads in infected cells in 384-well plates. Both treatments discriminate between RML prion infected CAD5 and non-infectious brain homogenate (NBH) treated CAD5 cells. **** = p-value < 0.0001, Student’s t-test **C:** The infectivity of PIPLC and PK-treated cell lysates was determined by infecting CAD5 cells with lysates as indicated. The signal intensity of the highest dilution was measured and compared to untreated RML CAD5 cell lysate. PIPLC-treated cells and PK-treated lysates showed similar infectivity titers. Prion-infected and NBH, as well as RML on *CAD5*^Δ*Prnp*^ were used for control. **D:** RML-infected CAD5 and GT-1/7 cells were transfected with non-targeting (NT) or *Prnp* targeting siRNAs in a 384-well plate and subjected to PK or PIPLC treatment. Z’-factors were calculated for each condition.

We then tested our assumption that PIPLC-resistant PrP (henceforth termed “PrP^PLC^”) is a plausible surrogate for prion infectivity. Therefore, we generated chronically RML6-prion infected CAD5 cells (RML CAD5) (Supplementary Fig. 1A) by treatment with mouse brain homogenate containing the Rocky Mountain Laboratory (RML) strain of prions (Avar et al., 2020; Solassol et al., 2003). For control, we used cells inoculated with non-infectious brain homogenate (NBH CAD5). We then measured PrP by QUIPPER and by PK digestion in 384-well microtiter plates. The readout yielded a clear separation between infected and non-infected cells (Fig. 1B). We conclude that QUIPPER can reliably detect prion infection.

Previous reports suggested that PIPLC treatment of chronically infected cells reduces the amount of PrP^Sc^ (Enari et al., 2001). In order to ensure that PrP^PLC^ can be used as a surrogate for prion infectivity, we performed a scrapie cell assay in endpoint format (SCEPA) (Mahal et al., 2008), arguably the most precise method to determine infectious prion titers *in cellula*. We inoculated naïve CAD5 cells with three decadic dilutions from lysates of RML-infected CAD5 cells treated with PIPLC just prior to lysis. For control, we used untreated RML CAD5 lysate and PK-treated lysate, as well as naïve and CAD5^Δ*pmp*^ cells inoculated with NBH or RML. After three passages, cells were spotted onto EliSPOT membranes and digested with PK to selectively detect infected cells (Fig. 1C). Image analysis of the optical density of the membranes showed that PrP^PLC^ retained full infectivity associated with prions.

We then compared the discriminatory power of QUIPPER vs. PK digestion for identifying modulators of prion propagation. Chronically infected RML CAD5 and RML GT-1/7 (Supplementary Fig. 1B) cells were treated with *Prnp*-targeting siRNAs or non-targeting (NT) siRNA controls in 384-well plates. Computation of the Z’-factor, a measure of the separation between positive and negative controls (Zhang et al., 1999), showed that QUIPPER outperformed PK digestion in both RML GT-1/7 and RML CAD5 cell lines (Fig. 1D). We opted to use RML GT-1/7 cells for the genome-wide screen because of their strong adherence to tissue culture plates, which facilitated their handling in 384-well microplates.

### Genome-wide screen for prion modifiers

We used a genome-wide murine siRNA library containing a pool of three distinct siRNAs per target transcript. Each siRNA mixture was dispensed in duplicate to a final concentration of 20 nM using an acoustic dispenser. Each 384-well plate was loaded with 264 gene-targeting siRNA triplets, 22 NT siRNA, and 22 *Prnp*-targeting siRNAs. The outermost wells were left blank as it was found to be prone to evaporation (Supplementary Fig. 1C). Controls and duplicates were strategically positioned for identifying and correcting any artifactual plate gradients, dispensing errors, or hotspots (Heinzer et al., 2021; Pease et al., 2019). Such gradients can arise from problems with the dispensing and aspiration steps or from temperature/humidity inhomogeneities during the incubation. After three days of culture, RealTime-Glo (RT-Glo), a reagent for cell-viability readout, and PIPLC were added and incubated at 37 °C for two hours. Subsequently, RT-Glo luminescence was measured, medium was aspirated, cells were lysed, and PrP^PLC^ was disaggregated and denatured. Finally, the antibody pairs were added to each well, and TR-FRET was measured after a 24 h incubation (Fig. 1A).

We screened a total of 136 plates entailing 17’582 in duplicates using siRNA triplets as well as 2’992 negative and positive controls, respectively. For each plate, heatmaps of TR-FRET and RT-Glo values (Supplementary Fig. 1D) were generated to detect any artifactual signal gradients or hotspots, which may have occurred during the screening process. 15’548 genes were assayed in duplicates and 2’030 genes in single measurements; four genes were not assayed during the primary screening. Z’-factors (Zhang et al., 1999) were >0.5 for 125 plates and 0 - 0.5 for 11 plates, confirming the robustness of the screen (Fig. 2A). A plot of all TR-FRET values obtained from the screen showed two non-overlapping populations corresponding to *Prnp*-targeting and NT controls, whereas the majority of the genes interrogated by the library had no effect on prion levels (Fig. 2B). The determination coefficient r^2^ between duplicates was 0.38, indicating a correlation sufficient to enable candidate selection (Taylor, 1990) (Fig. 2C).

**Figure 2.**
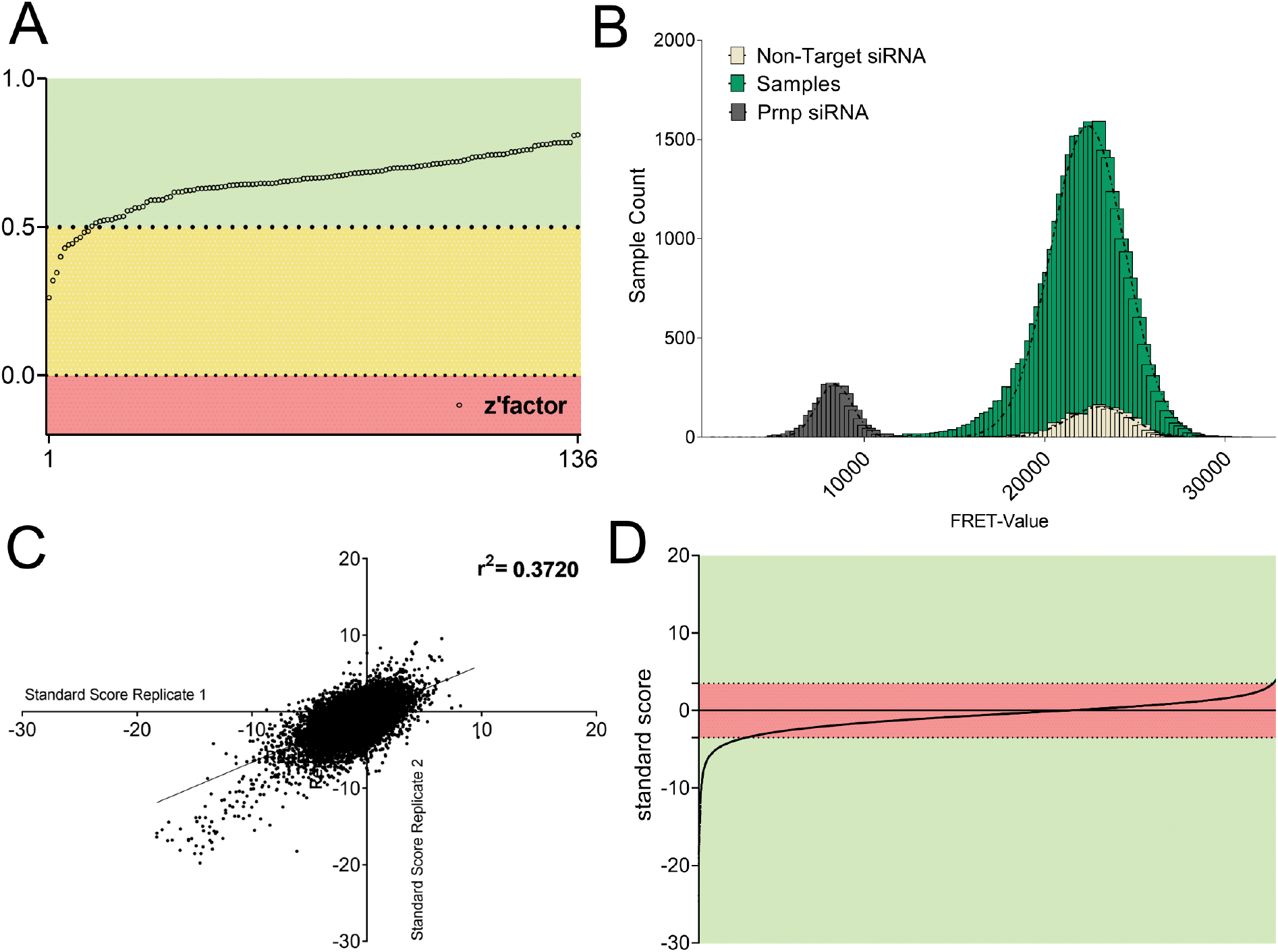
Whole-genome screen for prion modulators. **A:** Z’-factor for each plate of the whole genome screen representing the robustness of the screen based on the separability of the positive (*Prnp* targeting) and negative (nontargeting) controls. **B:** Histogram representing the influence of each protein-coding gene as well as NT and *Prnp* targeting controls, on prion levels. Abscissa: prion levels measured by FRET. Ordinate: number of genes yielding a given FRET-range. Controls showed a strong separation, allowing for confident hit-selection. Only few genes affected prion levels. **C:** Correlation of standard scores for all genes that were assayed in duplicates in the primary screen. r^2^: coefficient of determination. **D:** All individual data points from the primary screening. Genes reaching a z-score of [<3.5] ∪ [>3.5] in one or both duplicates were considered as hits (green area).

We then computed the standard score (z-score) for each candidate (Birmingham et al., 2009), and selected the top scoring 2’515 candidates (z-score=[<-3.5]∪[>3.5]) for a confirmatory screen (Fig. 2D). Of these candidates, 2’154 had a negative z-score, whereas only 361 genes had a positive z-score. Hence, 86% of modifiers, when suppressed, reduced PrP^PLC^ levels, whereas modifiers whose suppression enhanced PrP^PLC^ levels were rarer (14%) (for candidate selection process, also see Supplementary Fig. 1E).

### Confirmatory screens on prion modulators

Since PrP^C^ is necessary for prion propagation, some prion modifiers may act by changing PrP^C^ expression or localization, whereas others may act selectively on PrP^Sc^. We therefore performed two secondary screens. In the first screen, all top-scoring 2’515 hits were tested for the modulation of PrP^C^. To exclude any potential confounders, GT-1/7 were exposed to NBH and passaged identically to the prion-infected cells, and the assays were performed as in the previous screen except for the omission of PIPLC treatment. In a second screen, we performed QUIPPER on the same 2’515 hits and measured PrP^PLC^.

All plates passed quality control (Supplementary Fig. 2A) and the reproducibility of duplicates was high (r^2^ for the PrP^C^ and PrP^PLC^ subsets: 0.77 and 0.76 for QUIPPER and 0.8 and 0.88 for viability, respectively; Supplementary Fig. 2B and 2C). The TR-FRET scores of duplicates were averaged and a z-score for measuring the effect size of the manipulation of each gene was computed (Supplementary Table 1). The PrP^C^ and PrP^PLC^ subsets were strongly correlated (r^2^: 0.62), suggesting that selective PrP^PLC^ regulators are rare (Fig. 3A). Moreover, there was a strong correlation between the RT-Glo measurements for NBH and RML infected GT-1/7 cells (Supplementary Fig. 2D), implying that no gene knockdown resulted in synthetic lethality with prion infection under these conditions.

**Figure 3.**
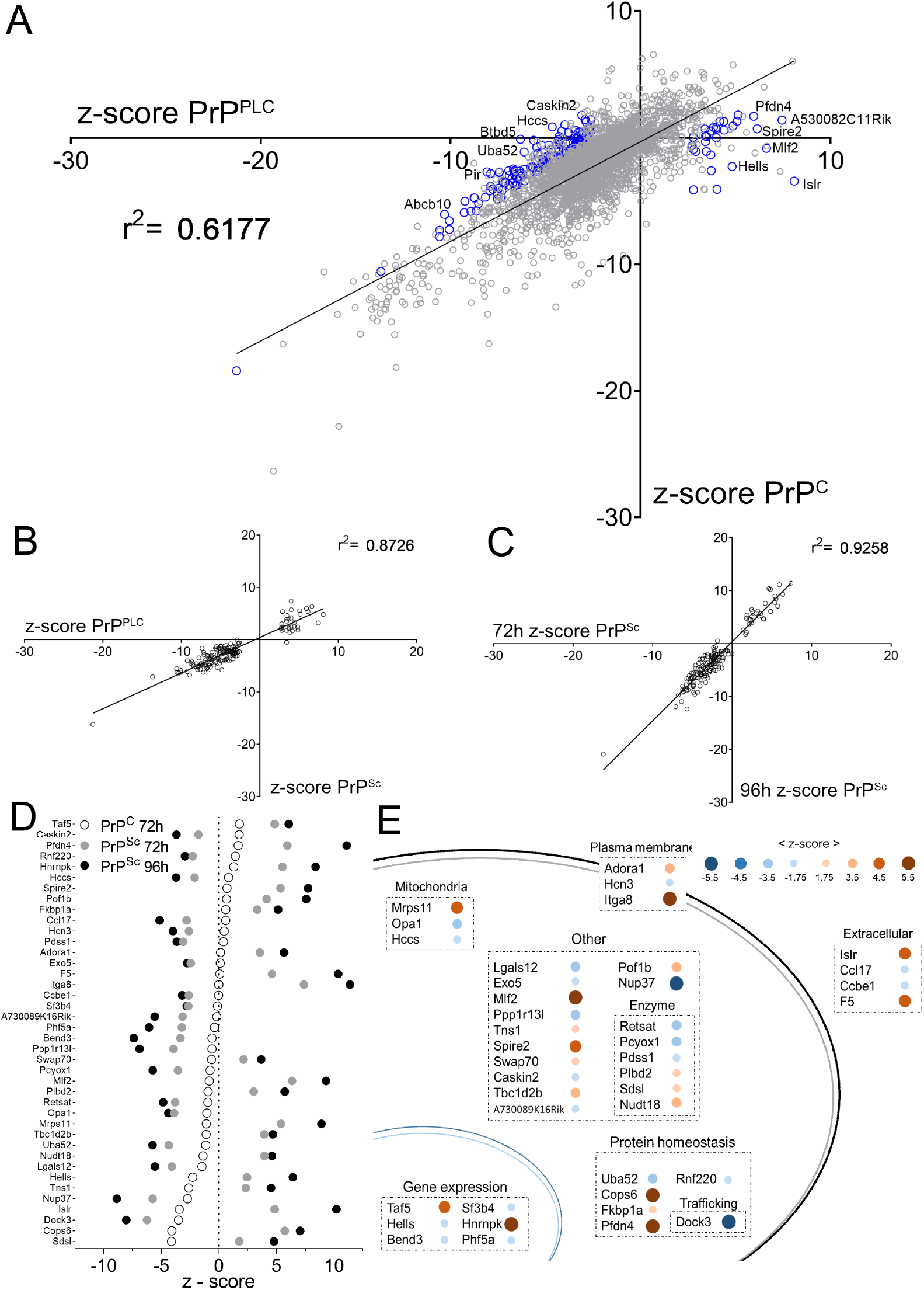
Secondary screens and shortlisting of 40 candidates. **A:** Regression of the values of the confirmatory screen for prion specific modulators in mock- and prion infected GT-1/7 cells. Z-scores of genes from the two independent screens, assessing either regulation of PrP^C^ or PrP^PLC^, yield a coefficient of determination (r^2^-value) of 0.62 indicating that most genes are modulating prion levels via regulating PrP^C^. Blue circles indicate 161 prion specific hits selected for downstream counter screens. The most conspicuous modulators were labelled. **B:** Correlation of z-scores obtained from a secondary screen to assess the effect of PK digestion on 161 PrP^PLC^ modulators in RML GT-1/7 cells after 72 hours of RNAi treatment. Z-scores of genes from the two independent screens (PIPLC or PK for two different sample preparation approaches) yield a coefficient of determination (r^2^) of 0.87, indicating that the candidates regulate PK-resistant prions. **C:** Correlation of the z-scores obtained from the counter-screenings to assess the effect of PK digestion on the prion modulators in RML GT-1/7 cells after 72 hours and 96 hours of RNAi treatment. The coefficient of determination (r^2^-value) of 0.93 and the increase in effect size for the prolonged treatment condition indicate a robust effect of the candidates on prion levels. **D:** Summary of the effect of the 40 shortlisted hits on PrP^C^ (after 72h) and PrP^Sc^ (after 72 and 96h), assayed using PK digestion, given as z-scores. **E:** Function and topology of the 40 hits. Blue dots: prion stabilizers; brown dots: prion limiters. Size and color saturation represent the effect size of each hit based on its z-score (72 hours RNAi treatment; PK readout).

We then applied layered criteria to identify genes modulating specifically PrP^PLC^. Firstly, in the repetition of the QUIPPER assay, only genes with a z-score [<-2.58] ∪ [>2.58], corresponding to a p value of 0.01, were considered hits. Secondly, in order to exclude any genes with a strong impact on viability, we limited the hit calling to samples in which the raw RT-Glo signal was not below 50% of the plate-specific NT control. Thirdly, we considered only genes whose effect size was ≥2.58 standard deviations higher for PrP^PLC^ than for PrP^C^ in uninfected cells. The latter criterion led to the exclusion of 99% of the overlap in two datasets. Lastly, all genes fulfilling these criteria were filtered according to their expression levels in RML GT-1/7 cells (Supplementary Table 1), thereby excluding genes that were not expressed (also see Supplementary Fig. 1E). This led to a list of 161 genes whose expression has a stabilizing (n=131) or limiting (n=30) effect on the amount of PrP^PL^C (Fig. 3A, blue circles).

### Role for prion specific modulators in sporadic Creutzfeldt-Jakob disease (sCJD) susceptibility

We wondered whether any of the 161 modulators of PrP^PLC^ may have a role in genetic susceptibility to sCJD (Heinzer et al., 2021). No gene passed the threshold for multiple hypothesis testing. The top ranked association was *DOCK3* (Multi-marker Analysis of GenoMic Annotation (MAGMA) unadjusted p=0.00063), a brain-resident guanine exchange factor serving as a binding partner to presenilin, which is implicated in Alzheimer’s disease and neurodegeneration (Bai et al., 2013; Chen et al., 2002; Chen et al., 2009; Tachi et al., 2012) (Supplementary Table 1).

### Validation of prion-specific regulators

Our observations suggest that QUIPPER may detect perturbations of prion propagation more sensitively than PK digestion. However, because QUIPPER is a new assay that has not yet been validated extensively by multiple laboratories, we subjected prion-infected GT-1/7 cells to PK digestion upon treatment with siRNA triplets corresponding to each of the 161 hits after 72 and 96 hours of siRNA treatment. We found a remarkable convergence between the QUIPPER and PK assays over the entire collection of genes (Fig. 3B), represented by the high r^2^ of 0.87, bolstering our confidence in the robustness of the targets identified. Prolonging the treatment with siRNA enhanced the effects observed (Fig. 3C). We then asked whether the hits were specific to a particular prion strain. For that, we infected GT-1/7 cells with the 22L strain of prions (Supplementary Fig. 3A) and treated them with siRNA triplets corresponding to 97 randomly selected hits, filling a 384-well plate. Most genes showed similar effects on RML and 22L-infected cells at both timepoints tested (Supplementary Fig. 3B).

In summary, 40 out of the 161 candidates showed a robust and consistent prion modulation across all detection methods (see Supplementary Figure 1E). 20 out of these 40 candidates reduce prion propagation upon silencing, and 20 candidates enhanced prion propagation, and henceforward are called stabilizers or limiters, respectively (Fig. 3D-E, Supplementary Table 1).

### Hnrnpk expression limits of prion propagation in mouse and human cells

Intriguingly, the suppression of *Hnrnpk,* an essential gene whose ablation causes cell death (Tsherniak et al., 2017), strongly enhanced prion levels while changing PrP^C^ levels only slightly. Hence, *Hnrnpk* acts as a limiter of prion propagation (Fig. 3D and Fig. 4A, left panel). We repeated these experiments in hovS, a human cell line expressing ovine but not human PrP^C^, which is readily infectible with the PG127 strain of ovine prions (Avar et al., 2020). Again, the downregulation of *HNRNPK* in prion-infected hovS cells led to an increase in PrP^Sc^ (Fig. 4A, right panel). Prion infection of hovS cells induces a prominent cytopathology consisting of cytosolic vacuolation. *HNRNPK* suppression exacerbated vacuolation, whereas ovine *PRNP* (*ovPRNP*) suppression completely abolished it, in line with the notion that prion levels determine the extent of hovS cytopathology (Supplementary Fig. 4A). To validate *HNRNPK* as a limiter of prion propagation independent of siRNA transfection, the experiment was additionally performed using shRNAs through lentiviral transduction in hovS. The results obtained with a shRNA targeting *HNRNPK* was congruent to the results obtained via siRNA transfection (Supplementary Fig. 4A and B), again highlighting the validity of *HNRNPK* as a modulator of prion formation.

**Figure 4.**
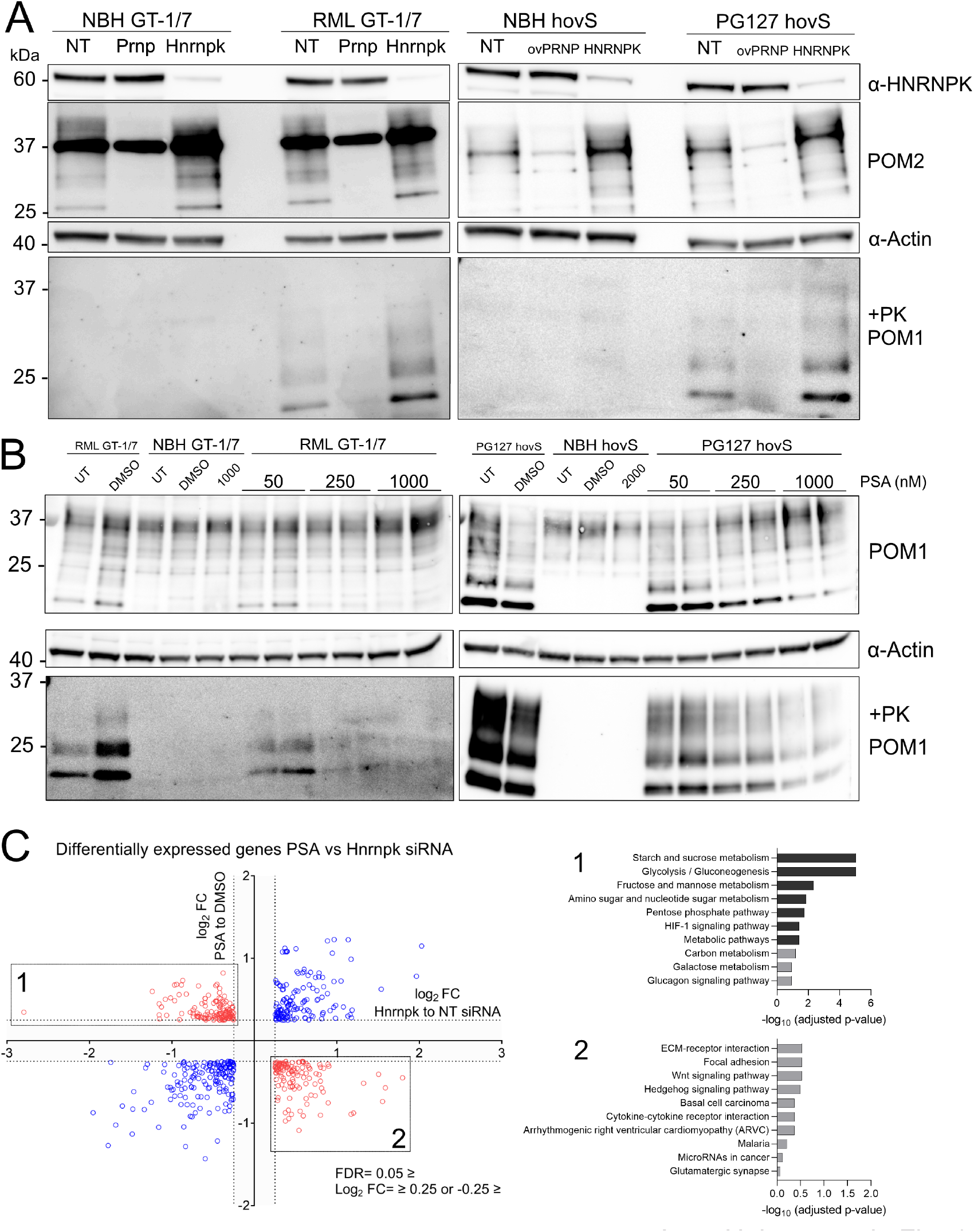
*Hnrnpk* limits prion levels in chronically infected mouse and human cell lines. **A:** Western blot showing *Hrnpk* siRNA transfection (96 hrs.) decreases Hrnpk protein levels while increases PrP^Sc^ in RML prion infected GT-1/7 and PG127 prion infected hovS cells. *Prnp* siRNAs suppressed both PrP^C^ and PrP^Sc^ as expected. α: anti **B:** Western Blot of PSA-treated uninfected and infected GT-1/7 and hovS cells. Increasing concentration of PSA leads to a more prominent reduction of PrP^Sc^ in mouse and human cells. **C:** Gene set overrepresentation analysis of differentially expressed genes (log_2_FC −0.25 ≥ or 0.25 ≤ and FDR ≤ 0.05) for siRNA mediated Hnrnpk downregulation or PSA treatment in RML GT-1/7 cells analyzed by RNAseq. Differentially regulated genes (up in siRNA treatment and down in PSA or vice versa) were overlapped and used for pathway analysis. No significantly enriched pathways are detected for upregulated genes in Hnrnpk and downregulated in PSA treatment. For the opposing direction, an enrichment of genes involved in glucose metabolism was detected.

To broaden our validation efforts, we treated cells with Psammaplysene A (PSA), which had been described to bind Hnrnpk (Boccitto et al., 2017). PSA treatment led to a dose-dependent decrease of PrP^Sc^ in prion-infected mouse and human cell lines (Fig. 4B) whereas PrP^C^ levels remain unaltered (Supplementary Fig. 4C). Next, we asked whether PSA indeed works on regulating PrP^Sc^ levels through its interaction with HNRNPK. As a knockout was not possible, due to the cell-essential nature of HNRNPK (Tsherniak et al., 2017) for long-term suppression, we used the shRNA constructs (HNRNPK-targeting and NT) and applied PSA at a concentration of 1 μM and started treatment 2 days after lentiviral transduction of shRNAs to allow time for HNRNPK downregulation to take place, and continued PSA treatment for 5 days. PK-digested western blots (Supplementary Fig. 4D) first confirmed that PSA does indeed reduce prion levels in combination with a NT shRNA construct and second, PSA’s effect does seem to be limited when HNRNPK shRNAs are applied. Moreover, we found that PSA does not alter HNRNPK levels (Supplementary Fig. 4D) and thus its antiprion effect potentially arises through enhancing the activity of HNRNPK.

We then performed RNAseq of RML GT-1/7 cells treated either with siRNAs against *Hnrnpk* or with PSA. As siRNA treatment increases prion levels and treatment with PSA leads to a decrease in prion levels, we intersected differentially expressed genes in opposing directions for both treatments. No significant enrichment was detected for genes that are upregulated in the siRNA treatment and downregulated in the PSA treatment. However, when genes that are downregulated in the siRNA treatment condition and upregulated in the PSA treatment condition were taken into account, genes related to glucose metabolism showed a significant enrichment (Fig. 4C).

### Psammaplysene A treatment leads to decrease in prion levels ex vivo and in vivo

We then asked whether a reduction of prions would be possible in a more physiological model system for prion infection and propagation. We found cerebellar organotypic cultured slice (COCS) (Falsig and Aguzzi, 2008) to be an ideal model to assess the efficacy of PSA treatment, as it represents a primary cell culture system with all relevant cell types represented as they are *in vivo*, while allowing easy experimental manipulation in a dish system. Therefore, we infected the cultures with prions using RML and started treatment of 1 μM PSA diluted in culture medium two weeks post-infection. Subsequently the COCS were homogenized and PK-resistant PrP^Sc^ amount was assessed with immunoblotting. We found that PSA treatment following prion infection significantly reduced prion levels (Fig. 5A), without altering PrP^C^ levels (Fig. 5B) further highlighting Hnrnpk as a limiter of prion propagation. Furthermore, we took advantage of a *Drosophila* model of prion propagation for the testing of *in vivo* efficacy of PSA. The model takes advantage of the ectopic overexpression of ovine PrP^C^ in *Drosophila melanogaster* (*Thackray et al., 2018*). Following prion infection, which is achieved through feeding the larvae with prion-containing food, flies produce bona fide prions and suffer from neurotoxicity associated with prions, which can be assessed with a negative geotaxis climbing assay. The experiment was performed as previously described (Thackray et al., 2018) and constant PSA treatment was achieved through feeding flies with the compound over the time course of 40 days. During this period, flies were subjected to a negative geotaxis climbing assay to assess whether locomotor deficits upon prion infection showed any improvement by treatment with PSA. Strikingly, a dose-dependent improvement of the performance of the animals was observed (Fig. 5C). Moreover, to address if the amelioration of the locomotor function corresponds to a reduction of prions, an RT-QuIC assay was performed to assess the amount of seeding active prions in the fly brains. Therefore, whole-head homogenates from 20 animals per treatment group were prepared and the samples were diluted 1:20 in PBS prior to their introduction into the RT-QuIC. Whereas flies treated with NBH and subsequently fed DMSO as controls yielded the same outcome as the prion-negative sample, an early peak in ThT signal were observed for flies that have been prion-infected and were later control fed using DMSO (Fig. 5D). Remarkably, a dose-dependent reduction of seeding active prions was observed in response to PSA treatment of flies infected with prions, as quantified based on the lag time (Shi et al., 2013) corresponding to the negative geotaxis climbing assay, demonstrating that PSA can alter prion levels *in vivo* in *Drosophila*.

**Figure 5.**
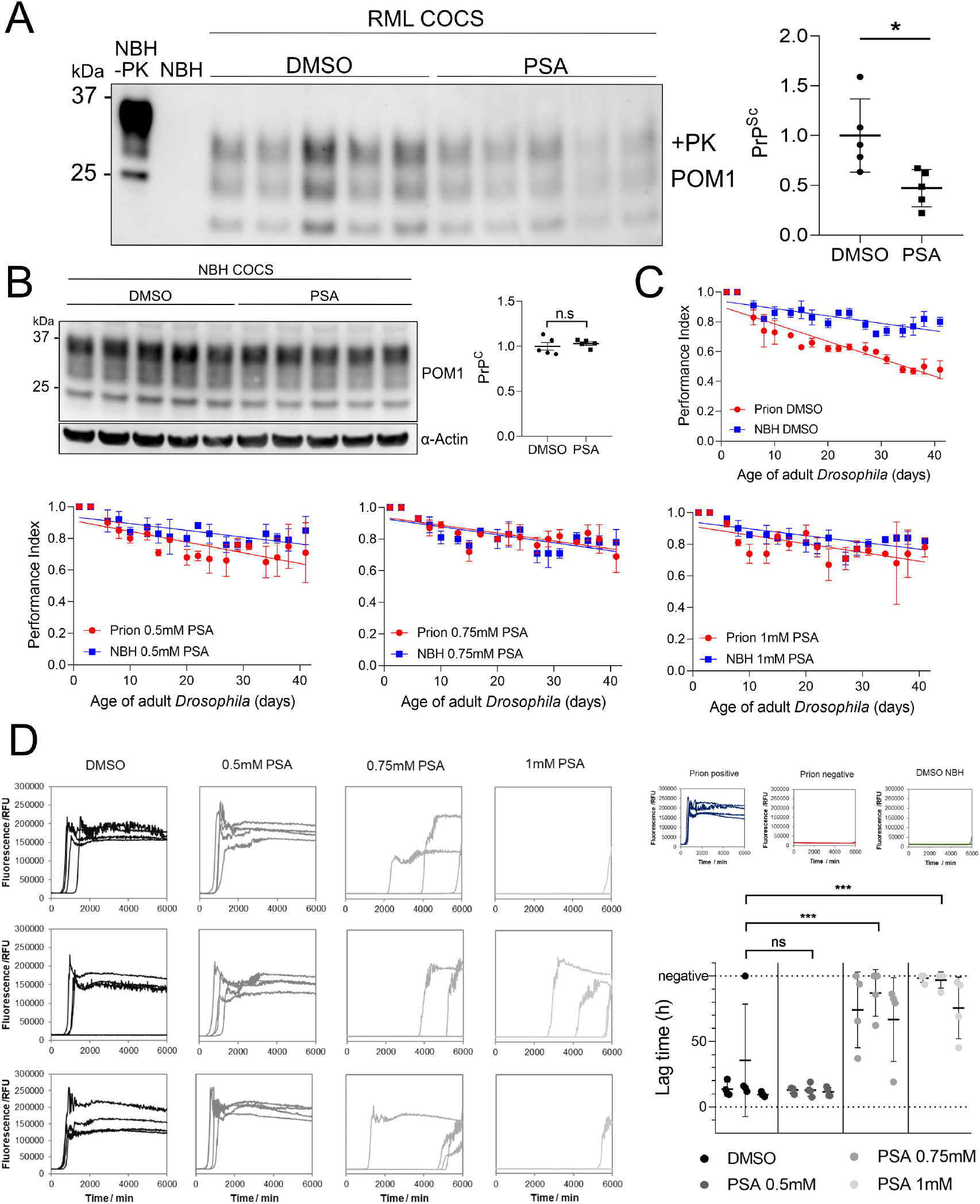
PSA reduces prion formation *ex vivo* and *in vivo*. **A:** Western Blot of RML prion infected COCS treated with 1 μM PSA or DMSO. PSA treatment was started two weeks after infection and continued until lysis. PSA reduced the amount of PrP^Sc^. Right panel: Quantification of the Western Blot; Values: mean ± SD. * p = 0 0211 (Student’s t-test). α: anti **B:** Western Blot analysis of NBH-treated COCS. PSA treatment was identical to the samples of A. PSA did not affect PrP^C^ expression. Right panel: Quantification of the Western Blot: Values represent mean ± SD. n.s. = not significant (student’s t-test). **C:** Negative geotaxis climbing assay in prion infected *Drosophila*. Flies were treated with DMSO, 0.5 mM PSA, 0.75 mM PSA, or 1 mM PSA at the larval stage and during adulthood for the duration of the assay. Climbing ability was assessed on groups of flies (n=3 x 15) three times a week and expressed as a performance index. Statistical analysis on the difference between PG127 prion infected versus control prion-free treatment group data in each graph was performed by an unpaired t test: DMSO: p = 0.0002; PSA: 0.5 mM: p = 0.0186; 0.7 5mM: p = n.s.; 1 mM: p = n.s. n.s. = not significant. **D:** RT-QuIC analysis of whole-head homogenates of prion infected *Drosophila*. For each sample, 10 male and 10 female heads from the same treatment group were homogenized, 1:20 diluted and applied to the RT-QuIC. Shown are the RT-QuIC reactions of three independent homogenates per treatment group, each assessed in quadruplicates. For quantification, the lag-time of each reaction was calculated and plotted in a graph. The assays were performed for 100 hours, samples not yielding a positive reaction are considered negative. *** = p ≤ 0.0006, n.s. = not significant, flies fed with NBH and treated with DMSO were used as a negative control, a standard prion-free and a prion-containing sample were used as assay controls.

## Discussion

Most previous attempts to identify prion modifiers were designed to test individual, biologically plausible candidate genes, which have been successful for example by uncovering the essential role of the B-cell receptor (Klein et al., 1997) and the CXCR5 chemokine receptor (Prinz et al., 2003) in prion propagation. Here, we attempted a radically different approach by performing a functional genomic screen to identify prion modifiers in cultured cells, with the only ideological constraint that all tested genes are proteincoding. Classically, prion detection relies on limited proteolysis, e.g. using proteinase K (Bendheim et al., 1984). As this approach is complex and imprecise, and therefore unsuitable to high-throughput screens, we have developed the QUIPPER, a robust method for the quantitative, sensitive, and scalable detection of prion propagation in cultured cells. The principle of QUIPPER relies on the finding that at steady state most PrP^C^ resides at the cell membrane and can be released by digesting with PIPLC, whereas prions remain associated to the cells (Borchelt et al., 1993; Stahl et al., 1990). We termed the identified PrP species PrP^PLC^ as it represents a mixture of prions and PrP^C^ from the endocytic compartments. To then allow the detection of PrP^PLC^, sodium hydroxide was added to induce disaggregation, enabling the recognition of the monomers by FRET donor-acceptor antibodies upon pH neutralization (Pease et al., 2019). Notably, the entire QUIPPER procedure is carried out as a one-pot assay, which greatly enhances its throughput and allows for extensive automation with programmable liquid handlers. Accordingly, QUIPPER enabled us to perform a whole-genome functional genomics screen using RNAi in chronically prion-infected mouse cells. Although we chose to employ QUIPPER in the context of a genetic screen, we believe that it will also be indispensable for future high-throughput drug discovery campaigns in search of antiprion compounds.

Following testing of the whole genome, a secondary screen consisting of 2515 candidates was performed and additionally assessed for changes in the levels of PrP^C^. Unsurprisingly, most of the primary candidates were regulating PrP^PLC^ by affecting the biosynthesis or degradation of PrP^C^ and PrP^PL^C specific regulators were rare. In addition, regulators of PrP^C^ demonstrated the strongest effects on prion levels, arguing for modulation of PrP^C^ as a potent and valid target in prion diseases (Raymond et al., 2019; Vallabh et al., 2020). Nevertheless, we selected 161 candidates showing an enhanced effect on PrP^PL^C and assessed those with classical PK digestion to identify modulators of PrP^Sc^, as some of the candidates might affect PrP^PL^C levels by changing the localization of PrP^C^, rendering it insensitive to PIPLC digestion. Thereby, we identified 40 hits that modulated prion levels, withstanding different biochemical prion detection methods. Among those, we reassuringly found hits that have been previously associated with prion disease (Marbiah et al., 2014; Wu et al., 2019), confirming the validity of our approach. When interrogated, these 40 hits did not reveal any significant associations with each other, such as working in concert on a known pathway or as interaction partners. However, it did not escape out attention that a group of hits have been previously associated with other neurodegenerative diseases (Bampton et al., 2021; Banerjee et al., 2017; Caminati and Procacci, 2020; Chen et al., 2001; Schludi et al., 2017; Stone et al., 2009; Takano et al., 2014), implying that there might be common host factors responsible for the progression of these ailments. One of them is Pfdn4, a member of the prefoldin complex (Takano et al., 2014), responsible for the folding of native peptides into their functional form. Although the occurrence of cytosolic PrP^C^ and its involvement in the formation of prions remains controversial (Ma and Lindquist, 2001; Ma and Lindquist, 2002; Ma et al., 2002), Prefoldin might be shielding cytosolic PrP^C^ from being accessed by prions, thereby preventing its incorporation, as upon downregulation of Pfdn4 we observed a strong increase in PrP^Sc^ levels. In addition, we identified several targets (Pof1b, Spire2, Swap70, Tns1, Dock3 and Itga8) that were independently described to be involved in actin binding or dynamics. Such a high prevalence might demonstrate the importance of the host-actin network in the cellular propagation of misfolded proteins. Indeed, actin has been shown to influence prion propagation in yeast (Dorweiler et al., 2020) and might be applicable as well in mammalian cells (See et al., 2021; Victoria and Zurzolo, 2017). Based on our data, the actin network might be a worthwhile target to further investigate. In addition, previous studies have shown promising results in modulating the propagation of pathological aggregates by interfering with the actin network (Rostami et al., 2017), in which actin may be a major component for cell-to-cell transmission of aggregates, through regulating processes such as synaptic vesicle exocytosis and others (Oliveira da Silva and Liz, 2020).

We put efforts forth in investigating Hnrnpk as a prion modulator, due to its strong effect size, ubiquitous expression (Uhlén et al., 2015), and its recent implications in several protein misfolding diseases (Bampton et al., 2021; Moujalled et al., 2015; Sidhu et al., 2022). As *Hnrnpk* is a cell-essential gene (Tsherniak et al., 2017), we followed several lines of validation in human and mouse cells as well as different modes of genetic perturbation. In all instances tested, *HNRNPK* downregulation led to an increase in prion levels, leading to the conclusion that HNRNPK is a host-factor regulating prion formation, irrespective of prion strain and even species in question. Moreover, in our human cell model for prion propagation, we found that the application of siRNAs and shRNAs targeting *HNRNPK*, led to a robust increase in prion-induced cytopathological vacuolation, and targeting PrP^C^ completely abolished it. In addition, a neuroprotective compound, Psammaplysene A, reported to bind Hnrnpk (Boccitto et al., 2017), allowed us to investigate the target pharmacologically. Psammaplysene A (PSA) application *in vitro, ex vivo* and *in vivo*, relieved prion propagation and ameliorated prion toxicity in a dose-dependent manner and therefore acts as a potent inhibitor of prion formation, repeatedly, independent of species in question and prion strain, in contrast to most other anti-prion drugs, which have been limited in their effect for different prion strains (Ghaemmaghami et al., 2010). The conclusions of this are twofold: First, it demonstrates the possibility and effectiveness of targeting a host-resident factor in prion diseases. Second, it suggests that different prion strains rely on the same host-factors for effective propagation, implying that host factors might provide efficient anti-prion targets independent of prion strains. Although the mode of action of PSA and its effect on Hnrnpk could so far not be clarified, we posit that Hnrnpk’s endogenous anti-prion function is enhanced upon PSA treatment, as we found no difference in Hnrnpk levels following PSA treatment. Since PSA only binds to Hnrnpk in the presence of RNA (Boccitto et al., 2017), therefore, potentially interfering with its RNA binding properties, RNA-binding of Hnrnpk might be crucial for limiting prion propagation by limiting the amount of free RNA which has been described as a potential scaffolding factor for the formation of prions (Deleault et al., 2003). Investigation of or mimicry of its function might provide a novel therapeutic approach for prion diseases and potentially other neurodegenerative diseases. Moreover, we are confident that other prion modifiers discovered in this study will be of great use to investigate cellular mechanisms of prion propagation, and perhaps represent general cellular host-factors of protein aggregation.

## Materials and Methods

### Cell culture

GT-1/7 cells (AccessionID: CVCL_0281, (Mellon et al., 1990), CAD5 cells (CATH-A-Differentiated, AccessionID: CVCL_0199, (Mahal et al., 2008), hovS cells (ovinized SH-SY5Y cells, produced in house) were cultured in T75 or T150 tissue culture flasks (TPP, Trasadingen, Switzerland) in OptiMEM containing Phenol unless stated otherwise (Gibco, Thermo Fisher Scientific, Waltham, MA, USA). As supplements 10% of FBS (Takata, Göteborg, Sweden), 1% of Non-Essential Amino Acids (NEAA, Gibco), 1% GlutaMaX (Gibco) and 1% of Penicillin/Streptomycin mix (P/S, Gibco) were used. During the screening process phenol was omitted from the media using a no-Phenol formulation for OptiMEM (Gibco), to eliminate a potential interference with the TR-FRET readout. During harvesting of the cells Accutase (Gibco) was used and collection of the detached cells was done using 1X phosphate buffered saline (PBS, Kantonsapotheke Zurich, Switzerland or Gibco) followed by centrifugation at 1400 rpm (Sorvall Legend XT, Thermo Fisher Scientific) for four minutes with the aim of eliminating dead cells. Later, cell counting was done using trypan blue (Gibco). For culturing hovS cells, 400 μg/mL geneticin (G418 sulfate, Life Technologies, Gibco) was added to the media. For freezing of the cell culture stocks, cells were resuspended in Bambanker Freezing Medium (LubioScience, Zurich, Switzerland) or DMSO (Sigma Aldrich, St. Louis, MO, USA) containing 10% FBS and stored in −80 °C or a liquid nitrogen tank, respectively.

### Prion infection of cells

For infection of cells with different strains of prions, a previously established protocol was followed (Avar et al., 2020). Cells were seeded in 6-well plates and the infection occurred for all prion strains (22L, RML, PG127) at the same weight/volume ratio of 0.25% brain homogenate containing prions or non-infectious brain homogenate (NBH) in a total culture media volume of 1.5 mL. Cells were incubated together with the infectious material for three days, followed by continuous splitting for eight passages to ensure a state of persistent prion infection was achieved.

### siRNA library preparation and screening

The whole genome Silencer Mouse siRNA library Version 3, consisting of three unique siRNAs targeting each annotated 17582 mouse gene (52746 siRNAs) was purchased from Thermo Fisher Scientific in a lyophilized format amounting to 0.25 nmol/siRNA. The library was resuspended, pooled and aliquoted in house as described (Heinzer et al., 2021) to a final concentration of 5 μM (1.67 μM each). For the screening procedure, the aliquoted pooled library was reformatted into the destination plates (384-well Cultur Plate, Perkin Elmer, Beaconsfield, UK) along with the *Prnp* targeting (Thermo Fisher Scientific) positive controls, NT controls (Thermo Fisher Scientific) serving as negative control and a cell deathinducting control (Qiagen, Hilden, Germany) to make up a final concentration of 20 nM. Plates were frozen at −40 °C until further use. On the assay day, plates were thawed and 5 μL of RNAiMAX (Invitrogen, Carlsbad, CA, USA) diluted in culture media without antibiotics (3% v/v) to a final concentration of 0.5% was dispensed with a Biotek MultiFlo FX multi-drop dispenser (Vinooski, VT, USA), followed by centrifugation (1000 rpm, 1 minute, Eppendorf 5804R, Hamburg, Germany) and subsequently the plates were incubated for 30 minutes at room temperature (RT). Later, cells (RML GT-1/7 or 22L GT-1/7, NBH GT-1/7, RML CAD5, NBH CAD5) were seeded on top of the siRNA-RNAiMAX mixture at a density of 12500 cells/well (3000 cells/well for RML CAD5 and NBH CAD5) in a total volume of 25 μL to achieve reverse transfection. Cells were incubated in a rotating tower incubator (LiCONiC StoreX STX, Schaanwald, Liechtenstein) for 70 hours and were removed for viability measurements. To measure the viability of cells, 10 μL of 4X concentrated Realtime-Glo (RT-Glo) reagent (Promega, Madison, WI, USA) diluted in antibiotic-free medium was dispensed. In addition, for the screening rounds involving a PIPLC readout, the enzyme was spiked into the media containing the viability reagent at a dilution of 1:800 and plates were returned to the rotating incubator for further incubation for another 2 hours. Subsequently, the luminescent signal was read out in an EnVision multimode plate reader (Perkin Elmer) at 37 °C. For the sample preparation with the PIPLC readout, culture media was aspirated from the plates using a Biotek EL406 plate washer and lysed with 13 μL of lysis buffer containing 0.5% Na-deoxycholate (Sigma Aldrich), 0.5% Triton-X (Sigma Aldrich) and EDTA-free cOmplete Mini protease inhibitor cocktail (Roche, Basel, Switzerland). The lysis of the contents of the wells was performed on a plate shaker (Eppendorf Thermo Mixer Comfort) for 10 minutes at 4 °C at 500 rpm shaking, followed by an additional incubation step at 4°C in still standing position for 1 hour. *B. cereus* PIPLC was produced in house as described previously (Hornemann et al., 2004; Ryan et al., 1996) in *E. coli*. For the PK readout, the protease inhibitor was omitted from the lysis buffer contents. Instead, the lysis buffer was spiked with a 2.5 μg/mL final concentration of PK, dispensed 10 μL, incubated for 30 minutes on a plate shaker at 37 °C with shaking, followed by addition of 3 μL of final 2.3 mM phenylmethylsulfonyl fluoride (PMSF, a serine hydrolase inhibitor) (Sigma Aldrich) diluted in PBS/isopropanol (Sigma Aldrich) mixture and incubated on a plate shaker for 10 minutes at RT and 500 rpm shaking conditions. For both the PIPLC and the PK readouts, the following steps involving denaturation and neutralization for sample preparation as well as the TR-FRET procedure for the final readout were identical. Denaturation was performed by dispensing 2 μL of 0.5M NaOH to a final concentration of 66.6 mM and plates were shaken with 500 rpm at RT for a total of 2 minutes followed by 8 minutes more of incubation time in still standing position. Denaturation of the fibrils was followed by a neutralization step to adjust the pH of the contents of the wells prior to the addition of the antibodies. 3 μL of 0.5 M NaH_2_PO_4_, was added to make up a final concentration of 83.3 mM and plates were shaken once more with 500 rpm at RT for a total of 2 minutes followed by 7 minutes more of incubation time in still standing position. For the read-out of PrP^C^ levels in the deconvolution screen, the usage of PIPLC, as well as the subsequent sample preparation steps were omitted. Instead, cells were lysed in 18 μL of lysis buffer. Finally, for all the approaches, 9 μL of TR-FRET antibody pairs, POM1 (Polymenidou et al., 2008) coupled to Allophycocyanin (APC) and POM19 coupled to Europium (EU) as previously described (Ballmer et al., 2017; Heinzer et al., 2021) were added to a final concentration of 5 and 2.5 nM, respectively. After each dispensing step during sample preparation and TR-FRET, plates were centrifuged at 1000 rpm for one minute. Plates were returned to the fridge for incubation overnight and read out on the following day on the EnVision (Perkin Elmer) plate reader with the previously described parameters (Ballmer et al., 2017).

### Data Analysis

Screening data was analyzed in the same way as previously described (Heinzer et al., 2021). Data quality and robustness was assessed at several steps during the analysis pipeline. Heatmaps were visually inspected for any occurrence of gradients, arising through dispensing errors. Wells with prominent dispensing errors were excluded from further analyses during the primary screen. Z’-Factor (Zhang et al., 1999; Zhang, 2011) was computed for each plate as a metric for separability of the positive (*Prnp* targeting siRNAs) and negative (NT siRNAs) controls. Net-FRET and z-score values for each candidate were calculated and regressed onto each other for assessing the reproducibility of the duplicates. Hit selection was based on the z-scores of each gene, as explained in the results section of this manuscript. Results were depicted using GraphPad Prism.

### RNA-Seq experiments

RNeasy Mini Kit (Qiagen) was used for RNA-extraction according to the manufacturer’s guidelines. Libraries were prepped with the Illumina TruSeq stranded mRNA protocol (Illumina, San Diego, CA, USA) and quality control (QC) was assessed on the Agilent 4200 TapeStation System (Agilent Technologies, Santa Clara, CA, USA). Subsequently, libraries were pooled equimolecular and sequenced on the Illumina NovaSeq6000 platform with single-end 100 bp reads. Sequencing depth was around 20 million reads per sample. Experiments were run in biological duplicates, unless otherwise stated. Data analysis was done using a previously established pipeline (Hatakeyama et al., 2016) (Supplementary Table 1). Pathway analysis for significantly enriched cellular processes or components were done using WEB-based GEne SeT AnaLysis Toolkit (WebGestalt) (Wang et al., 2017) using the KEGG functional category version 88.2 dated 11/01/2018.

### siRNA transfections and PSA treatment in large-well format

Cells were seeded at a density of 500’000 cells/well for RML GT-1/7 and 450’000 cells/well for PG127 hovS in 1.5 mL of culture medium in 6-well plates (TPP). Next day, media was exchanged with optiMEM (Gibco) with no antibiotics. To enable efficient transfection, RNAiMAX (Thermo Fisher Scientific) with the same concentration as in the screening process was added (final conc. for hovS: 0.3%). Next day, siRNAs diluted in water were mixed with RNAiMAX and a final concentration of 20 μM (10 μM for hovS) was administered to the culture media in a dropwise manner to achieve forward transfection. Incubation lasted 72 or 96 hours, as indicated per experiment. Media containing siRNAs was aspirated and cells were washed once with 1X PBS (Kantonsapotheke) and lysed for downstream analysis. Imaging of the cells was performed using a Nikon T2 Eclipse (Nikon, Tokyo, Japan) microscope and images were processed using ImageJ (Schneider et al., 2012). PSA (Aobious Inc., Gloucester, MA, USA) was freshly prepared at the beginning of any experiment by serial dilution in DMSO. For each concentration, one aliquot per day of treatment was stored at −20°C. The desired final concentrations of PSA were prepared daily by diluting 1 to 1000 the DMSO stock aliquot in culture media. In the PSA experiments, the cells were seeded in a total volume of 1.5 mL media in six-well plates at a density of 300’000 PG127 and NBH hovS cells 450’000 and RML and NBH GT-1/7 cells/well. From the following day for the next 5 days, daily medium changes were performed with freshly prepared media at differing concentrations of PSA.

### Immunoblotting

Lysed cells were centrifuged at 2’000X G for 5 minutes. BCA assay (Pierce, Thermo Fisher Scientific) was used to measure the total protein content of each sample and for all downstream analysis involving proteins, sample volume was adapted to contain the same amount. For immunoblotting, samples were prepared, diluted to achieve the same total protein concentration, and digested using PK (final concentration 2.5 μg/mL). Digestion was stopped with boiling the samples after addition of 1mM final Dithiothreitol (DTT, Bio-Rad, Hercules, CA, USA) in NuPAGE 4X LDS loading buffer (Thermo Fisher Scientific). Samples were then loaded onto a NuPAGE 4-12% Bis-Tris gradient gel (Invitrogen, Thermo Fisher Scientific) and blotted onto a nitrocellulose membrane using the iBlot dry transfer system (Invitrogen, Thermo Fisher Scientific). Membrane was blocked using 5% SureBlock (LubioScience) diluted in 1X PBS containing 0.1% Tween-20 (PBST, Sigma Aldrich) for 30 minutes. Membranes were then incubated with primary antibodies diluted in 1% SureBlock-PBST (POM1, POM2 as anti-PrP antibody, 300 ng/mL final concentration (Polymenidou et al., 2008), anti-Hnrnpk antibody (ab70492, 1:5000 diluted, Abcam, Cambridge, UK) overnight at 4 °C under shaking conditions. For detection, antimouse HRP or anti-rabbit HRP (Bio-Rad) was diluted 1:5000 in 1% SureBlock-PBST. Imaging was done on LAS-3000 System (Fujifilm, Tokyo, Japan).

### Cerebellar slice culture experiments

Animal experiments were performed in accordance with the Swiss Animal Protection law and under a permit issued by the Canton of Zurich (Nr: 236/2019). Cerebellar slices were prepared from 12-day-old *Tg*a20 mouse pups according to our published protocol (Falsig and Aguzzi, 2008). Briefly, acutely dissected cerebella were embedded in 2% low melting point agarose and cut into 350-μm thick sections with a Leica vibratome in ice-cold Gey’s balanced salt solution (GBSS) supplemented with kynurenic acid and glucose. For prion infection, slices were exposed to 0.001% brain homogenate derived from terminally sick RML6 prion inoculated mice (or normal brain homogenate as control) for 1 hour at 4 °C on a shaker. After several washes, six to eight slices were put onto a Millicell-CM Biopore PTFE membrane insert (Merck Millipore) and cultured in an incubator on top of slice culture medium containing 50% vol/vol MEM, 25% vol/vol basal medium Eagle, 25% vol/vol horse serum, 0.65% w/vol glucose, 1% vol/vol penicillin/streptomycin and 1% Glutamax. Culture medium was changed three times per week. To treat the cultured slices with PSA, stock PSA solution prepared in DMSO was diluted into the culture medium with a final concentration of 1 μM. Culture medium with the same concentration of DMSO was used as control. Fresh PSA was supplied to the media and media was changed every two to three days for a total duration of one month. At the end of the experiments, cultured slices were collected into the RIPA buffer and homogenized. Protein concentrations in the lysates were quantified using the BCA method. To detect prions by western blotting, lysates containing 50 μg total proteins were mixed with proteinase K (with a final concentration of 20 μg/mL) in a reaction volume of 30 μL and incubated at 37 °C for 30 minutes. After mixing each digested sample with 10 μl 4X loading buffer and boiled at 95 °C for 5 minutes, 18 μL of each sample was loaded onto the gel for western blotting. Quantification was done using ImageJ software and results were plotted using GraphPad Prism8. To assess statistical significance an unpaired Student’s t-test was performed.

### SCEPA for detection of infectivity in PK and PIPLC treated cells

For assessing whether prion infectivity is intact after treatment with PK (Roche) or PIPLC (Thermo Fisher Scientific) 100.000 RML CAD5 cells were seeded in 2 mL of full culture media and grown to confluency for four days. Later, samples for PIPLC were treated with 0.1U/mL of the enzyme which was spiked into the media and placed back in the incubator for one hour. Cells were then harvested in 1X PBS, centrifuged at 1500 XG for 5 minutes for pelleting and finally the PBS was aspirated. 40 μL 1X PBS was added to the cell pellet and lysis of the contents of the tube was followed with 5 freeze-thaw cycles in liquid nitrogen with vortexing. Subsequently, the lysate was centrifuged at 1000XG for three minutes and supernatant amounting to 50 μL was transferred to a new Eppendorf tube. For PK digestion, 10 μL of 15 μg/mL PK was added to the cells and digestion followed for 30 minutes at 37 °C. Digestion was stopped using 5 μL of PMSF (Sigma Aldrich) (final concentration 2.3 mM) under the same conditions described in the screening section of the manuscript. Untreated lysates as well as PIPLC treated samples were harvested as described above and 15 μL PBS was added to the samples to make up the same volume. SCEPA was performed as previously described (Sorce et al., 2020).

### shRNA design, cloning, lentiviral vector preparation and transduction

Using NEBuilder HiFi DNA Assembly Cloning Kit (NEB #E5520), eGFP coding sequence (Addgene # 17397) was cloned into our Auto-TDP-43-HA LV transfer vector (Hruska-Plochan et al., 2021), replacing TDP-43-HA, making Auto-EGFP. Custom MHP_shRNA cassette was synthetized by GenScript so that human U6 promoter is directly followed by random sequence of 54bp flanked by HindIII and PacI restriction sites. TTTTTTT was used for efficient PolIII termination (Gao et al., 2018) and TGTGCTT for loop (Jensen et al., 2012). This cassette was then cloned into Auto-EGFP via HiFi assembly, upstream from TRE promoter, generating pSHE LV transfer vector.

HNRNPK transcript variant 5 (NM_001318186.2) was then used as an input sequence for shRNA design using the following algorithms: Broad Institute GPP Web Portal, Invitrogen BLOCK-iT™ RNAi Designer, Kay Lab siRNA/shRNA/Oligo Optimal Design tool and RNAinverse server of ViennaRNA Web Services. 5 highest-scoring target sequences per algorithm were kept and 2 final target sequences were selected using rational design following the standard rules for pre-miRNA-like shRNA design (Bofill-De Ros and Gu, 2016), including Dicer loop-counting rule (Gu et al., 2012). Q5 site directed mutagenesis (NEB #E0554S) was used to clone the designed shRNAs into pSHE, substituting the random sequence of the original cassette, generating pSHE-shHNRNPKa and pSHE-shHNRNPKb LV transfer vectors. shRNA targeting the HaloTag sequence was then designed as a non-targeting shRNA control and was cloned into pSHE as decribed above, generating pSHE-shHaloTag. pSHE vectors were then packaged into lentivirus (LV) as described previously (Hruska-Plochan et al., 2021). The resulting lentiviral pellets were then resuspended in PBS to achieve 10x concentrated LV preparations, which were titrated using Lenti-X™ GoStix™ Plus (Takara #631280). 10x concentrate of pSHE LVs was then used at 300 ng (of lentiviral p24 protein as per GoStix Value (GV)) per well of a 6 well plate of hovS cells pipetting the LV concentrate directly onto the culture (dropwise). Spent hovS media was then added to reach 1000μl total. Medium was exchanged completely the following day.

HNRNPK knockdown efficacy of pSHE-shHNRNPKa and pSHE-shHNRNPKb was assessed in SH- SY5Y cells and pSHE-shHNRNPKa was selected for hovS experiments. For immunoblotting of transduced cells, see above.

Primer list: shHNRNPKa_F 5’-TGCTTAAGCATTCCACAGCATCTTTTTTTAATTAACATGGTCCCAGC-3’
shHNRNPKa_R 5’-AGCACAGCTTAAGCATTCCACAGCATCAAGCTTTCGTCCTTTCCAC-3’
shHNRNPKb_F 5’-TGCTTAAACCACCAACAATAACTTTTTTTAATTAACATGGTCCCAGC-3’
shHNRNPKb_R 5’-AGCACAGCTTAAACCACCAACAATAACAAGCTTTCGTCCTTTCCAC-3’
shHaloTag_F 5’-TGCTAAATGCAATACCTTTGACTTTTTTTAATTAACATGGTCCCAGC-3’
shHaloTag_R 5’-AGCACAGCTAAATGCAATACCTTTGACAAGCTTTCGTCCTTTCCAC-3’

### RT-QuIC assay

The reaction buffer of the RT-QuIC consisted of 1 mM EDTA (Life Technologies), 10 μM thioflavin T, 170 mM NaCl, and 1× PBS (incl. 130 mM NaCl) and HaPrP23-231 filtered using 100-kD centrifugal filters (Pall Nanosep OD100C34) at a concentration of 0.1 mg/ml. Fly brain homogenates were diluted in PBS and 2 μl of the diluted homogenates were used to assess seeding activity, resuspended in 98ul of assay buffer in a 96-well plate format. The plate was loaded into a FLUOstar Omega plate reader (BMG Labtech) and the shaking cycles were set as follows: 7× (90 s shaking; 900 rpm [double orbital]; 30 s rest) and 60 s reading. Reading was carried out with excitation at 450 nm and emission at 480 nm every 15 min. The amplification was performed at 42°C for 105 h. Four replicates per sample were measured. Lag time was determined as timepoint at which the sample reached 20’000 RFU.

### Fly stocks

The *UAS*-PrP fly line w; M{VRQ-PrP(GPI), 3xP3-RFP.attP}ZH-51D transgenic for V^136^R^154^Q^171^ (VRQ) ovine PrP, expressed with an N-terminal leader peptide and C-terminal GPI signal sequence, was generated by PhiC31 site-specific transformation (Thackray et al., 2012) with *Cre*-mediated removal of the red fluorescent protein (RFP) as described previously (Thackray et al., 2014). The Elav-GAL4(P{w[+mW.hs]=GawB}elav[C155]) pan neuronal driver fly line was obtained from the Department of Genetics, University of Cambridge, UK. *Drosophila* were raised on standard cornmeal media at 25 °C and maintained at low to medium density.

### PSA treatment and prion inoculation of Drosophila

PSA was prepared in DMSO to give 0 mM, 0.5 mM, 0.75 mM or 1 mM final concentration of drug. Two hundred microlitres of the relevant dilution of PSA or DMSO alone were added to the top of fly feed in 3-inch plastic vials and the feed allowed to dry for 24 hours prior to use. A cross between the *UAS*-PrP fly line and the Elav-GAL4 driver fly line was set up in each of the drug-treated or control fly food vials described above. The parental flies were removed from the vials once first instar larvae were evident. For prion inoculation, 250 μL of 1% (w/v) PG127 scrapie-positive sheep brain homogenate (Andréoletti et al., 2011)prepared in PBS pH 7.4, were added to drug-treated third instar VRQ ovine PrP *Drosophila* larvae. Following eclosion (i.e. hatching), flies were collected that were transgenic for VRQ ovine PrP expressed pan neuronally and were transferred to fresh drug-treated vials every other day for the duration of the study (40 days). The locomotor ability of flies was assessed in a negative geotaxis climbing assay as previously described (Thackray et al., 2018). *Drosophila* head homogenate was prepared from 5 and 40 day old flies as previously described (Thackray et al., 2018) and subjected to RT-QuIC analysis as described in the materials and methods. Statistical analysis of the negative geotaxis climbing assay data was performed by the unpaired *t*-test, using Prism (GraphPad Software Inc, San Diego, USA).

---

**Supplementary Figure 1.**
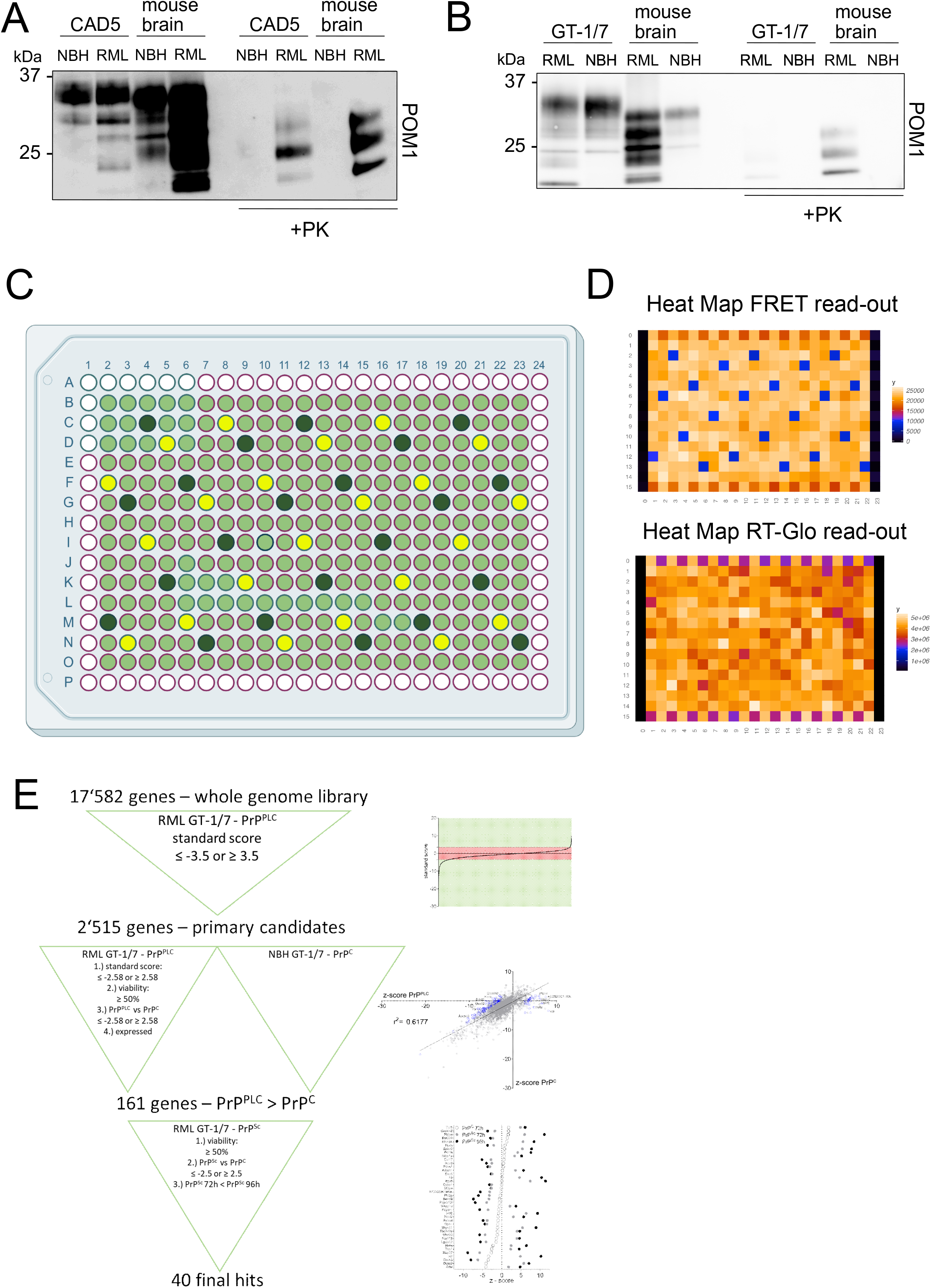
QUIPPER assay, Example plate maps **A)** Western Blot analysis of chronically RML6-infected CAD5 cells following PK digestion. Anti-PrP antibody POM1 is used for probing the membrane. Brain homogenates were used as controls. **B)** Western Blot analysis of chronically RML6-infected GT-1/7 cells after PK treatment. Membrane is probed with anti-PrP antibody POM1. Brain homogenates were used as controls. **C)** Plate map used in the screen depicting controls and samples. Light green represents wells containing siRNAs from the library, dark green represents wells containing Prnp-targeting control siRNAs and yellow represents wells containing non-targeting control siRNAs. **D)** Examples of heat maps for FRET as well as viability read-out from the primary screen. **E)** Schematic of the hit-selection process over all the screens performed in this study with the corresponding criteria.

**Supplementary Figure 2.**
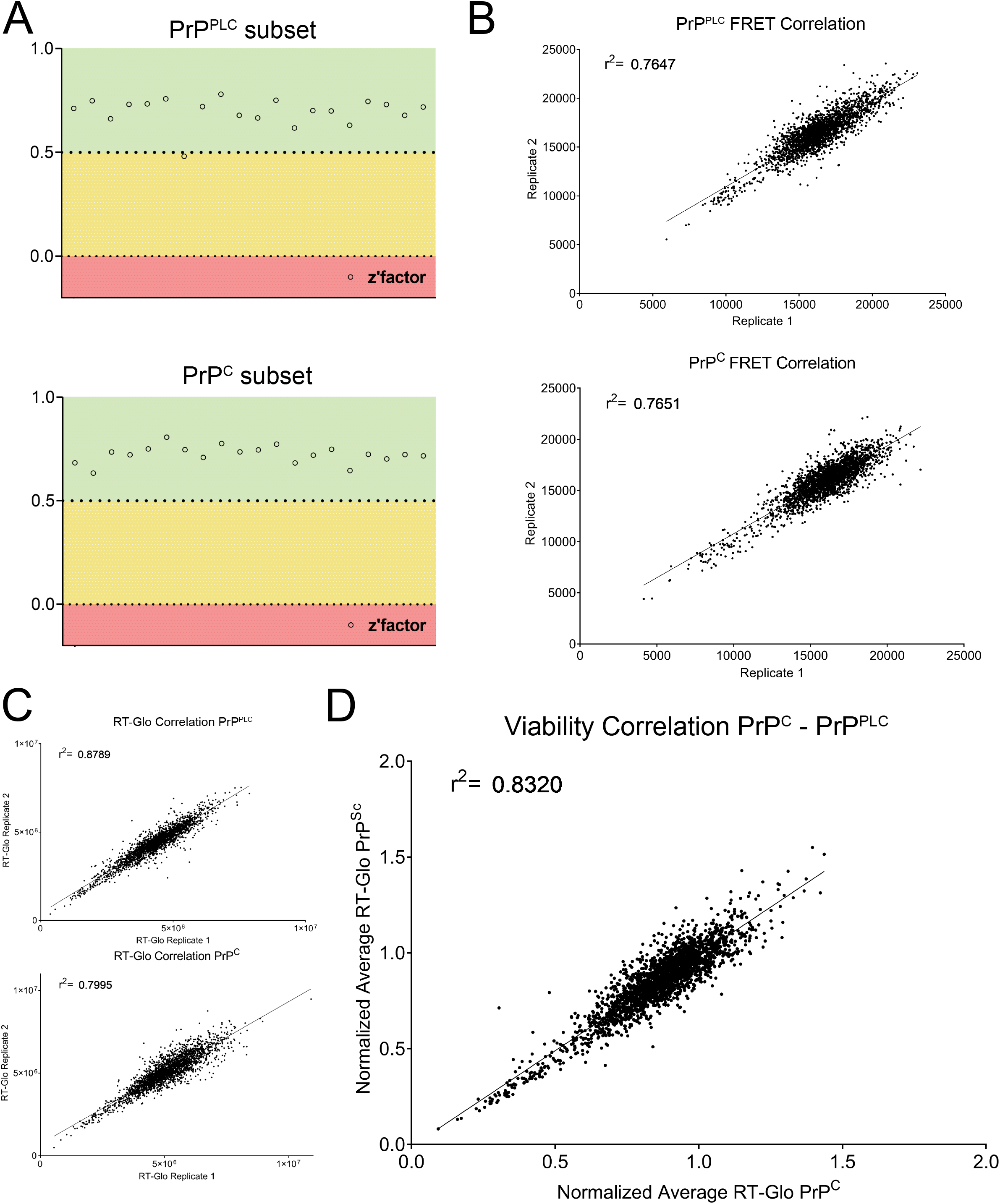
Secondary screen quality metrics and viability readout **A)** Z’-factor for each plate of both deconvolution screens (for regulators of PrP^PLC^, top panel, for regulators of PrP^C^, bottom panel) representing the robustness of the screens based on the separability of the positive (*Prnp* targeting) and negative (non-targeting) control **B)** Duplicate correlation of FRET-data for each screen. Coefficient of determination (r^2^-value) is depicted in the graph. **C)** Duplicate correlation of viability-data measured using RealTime-Glo™ luminescence readout for each screen. Coefficient of determination (r^2^-value) is depicted in the graph. **D)** Duplicates from C were averaged and normalized for each screen, and the two normalized values for each gene is correlated to assess if any gene regulates viability dependent on prion infection. The high r^2^-value, as well as the lack of outliers demonstrates the lack of synthetic lethal genes in the subset assessed in the deconvolution screen.

**Supplementary Figure 3.**
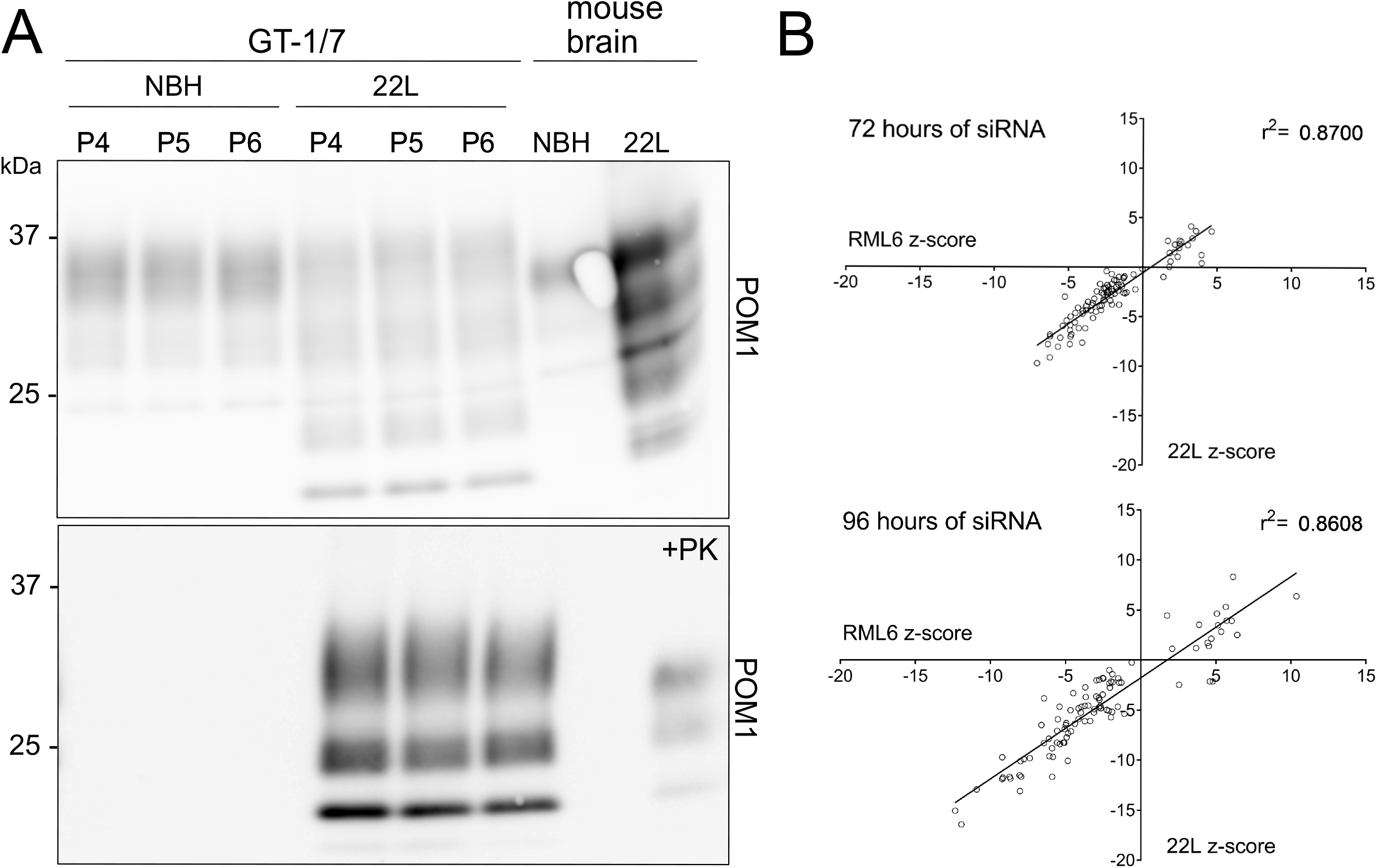
Prion strain dependence of the 97 candidates **A)** Western Blot analysis of chronically 22L-infected GT-1/7 cells after PK treatment. Membrane is probed with anti-PrP antibody POM1. P4, P5 and P6 corresponds to the amount of passaging after exposure to 22L brain homogenate. Brain homogenates were used as controls. **B)** Correlation of the effect of 97 candidates on their effect on PrP^Sc^ of two different prion strains (RML6 and 22L). Next to RML6 infected GT-1/7, 97 hits were assessed for their effect on 22L prion infected GT-1/7 for 72 hours as well as 96 hour-long treatment duration. The high coefficient of determination (r^2^-value) of the effect observed for both prion strains indicates that the candidates do not show strain-specificity.

**Supplementary Figure 4.**
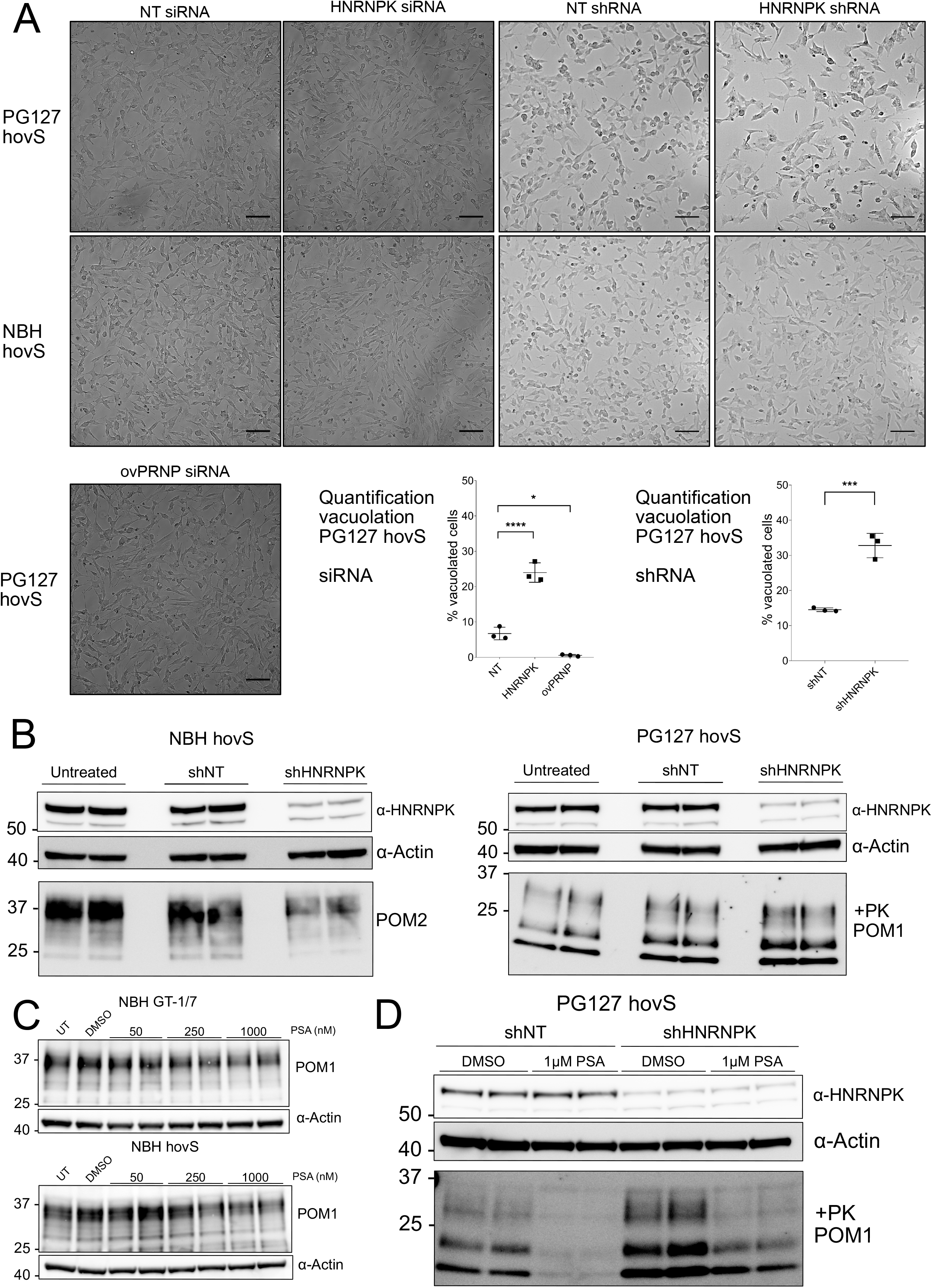
Phenotypic response of PG127prion infected hovS upon *HNRNPK* downregulation using siRNA or shRNA **A)** Brightfield microscopy images of the effect of siRNA and shRNA mediated *HNRNPK* downregulation on prion-induced vacuolation in PG127 hovS cells. The siRNA and shRNA mediated downregulation of *HNRNPK* in PG127 hovS leads to an enhanced cytopathological vacuolation phenotype when compared to NT siRNA or shRNA, corresponding to the increased level of prions seen in Fig. 4A and Supplementary Figure 4B. ov*PRNP* siRNA transfected, as well as uninfected cells were used as controls. Downregulation of ov*PRNP* in the infected hovS eliminates the vacuoles; *HNRNPK* downregulation in uninfected cells does not yield a vacuolation phenotype. Lower left panel: Quantification of vacuoles of NT, *HNRNPK* and *ovPRNP* siRNA treated PG127 hovS. Lower right panel: Quantification of vacuoles of NT and *HNRNPK* shRNA treated PG127 hovS. Cells from pictures at three different positions in the well were manually counted and the amount of vacuolated cells was normalized to the total amount of cells. Values represent mean ± SD. * p = 0.0113, *** p = 0.0008, **** p = <0.0001 (Dunnett’s multiple comparisons test). Scale bar = 100 μm **B)** Western blot showing *HNRNPK* shRNA transduction (96 hrs.) decreases HNRNPK protein levels while increases PrP^Sc^ in PG127 prion infected hovS cells. *α*: anti. **C)** Western Blot analysis of the effect of PSA treatment on PrP^C^ levels in uninfected GT-1/7 as well as hovS cells. PSA treatment does not affect the levels of PrP^C^ when compared to untreated or DMSO treated cells. **D)** Western Blot analysis of the effect of PSA treatment on PrP^Sc^ and HNRNPK levels in PG127 prion infected hovS cells following transduction of control (NT) or HNRNPK targeting shRNAs. PSA treatment does not affect the levels of HNRNPK (upper panel) when compared to DMSO treated cells. The antiprion effect of PSA is limited upon silencing of HNRNPK (lower panel). α: anti.

## Supplementary Table

**Tab 1:** Z-scores and normalized viability values for each gene assessed in the deconvolution screen

**Tab 2:** RNA-Seq of RML prion infected GT-1/7

**Tab 3:** MAGMA and VEGAS scores for comparing the 161 candidates for which human orthologues are available to a sCJD GWAS dataset.

**Tab 4:** Z-scores and normalized viability values for the 40 shortlisted genes assessed in the PrP^S^C screen, using PK readout after 72 hours and 96 hours of treatment, as well as their corresponding gene symbols for *Homo sapiens*.

**Tab 5.1:** RNAseq data of *Hnrnpk* downregulated RML GT-1/7 cells

**Tab 5.2:** RNAseq data of 1 μM PSA treated RML GT-1/7 cells

## Acknowledgements

AA is the recipient of an Advanced Grant of the European Research Council and grants from the Swiss National Research Foundation, the Nomis Foundation, the Swiss Personalized Health Network (SPHN, 2017DRI17), and a donation from the estate of Dr. Hans Salvisberg. We would like to thank Dr. Emilio Yangüez and Dr. Maria Domenica Moccia and the Functional Genomics Center Zurich (FGCZ) for their help with the RNA-Sequencing experiment. We thank Dr. M. Ryan and Dr. O.H. Griffith for providing us with the expression system for *B. cereus* PIPLC. We thank Prof. Simon Mead for help with the analysis of the sCJD dataset. We thank Prof. Robert Kalb and Dr. Marco Boccitto for insightful discussions. In addition, we wholeheartedly thank Irina Abakumova and Rita Moos for their technical help. Summary Fig. 1A depicting the QUIPPER assay and Supplementary Fig. 1C have been created with BioRender.

## Author contributions

Conceived and designed the experiments: MA, DH, AA. Supervised the study: SH, MP, RB, AA. Contributions to experimental work; siRNA primary and counter-screens: MA, DH, siRNA reformatting: MA, DH, DP, ME, bioinformatics: MA, DH, ES, AC, PSA experiments: MA, DH, SS, proposed the QUIPPER assay: AKKL, PIPLC purification: SH, shRNA design, production and validation: MHP, shRNA experiments in hovS: MA, DH, RNA-Seq experiment: MA, DH, COCS experiments and data analysis: YL, provided reagents and experimental help: JAY, ML. *Drosophila melanogaster* experiments and data analysis: MA, DH, AT, RB. Wrote the paper: MA, DH, AA.

